# Translation as a Biosignature

**DOI:** 10.1101/2023.08.10.552839

**Authors:** Jordan M. McKaig, MinGyu Kim, Christopher E. Carr

**Author notes:** Correspondence to. Address: ESM Building, Room G10, 620 Cherry St NW, Atlanta, GA 30332, USA.

## Abstract

Life on Earth relies on mechanisms to store heritable information and translate this information into cellular machinery required for biological activity. In all known life, storage, regulation, and translation are provided by DNA, RNA, and ribosomes. Life beyond Earth, even if ancestrally or chemically distinct from life *as we know it* may utilize similar structures: it has been proposed that charged linear polymers analogous to nucleic acids may be responsible for storage and regulation of genetic information in non-terran biochemical systems. We further propose that a ribosome-like structure may also exist in such a system, due to the evolutionary advantages of separating heritability from cellular machinery. Here, we use a solid-state nanopore to detect DNA, RNA, and ribosomes, and demonstrate that machine learning can distinguish between biomolecule samples and accurately classify new data. This work is intended to serve as a proof of principal that such biosignatures (i.e., informational polymers or translation apparatuses) could be detected, for example, as part of future missions targeting extant life on Ocean Worlds. A negative detection does not imply the absence of life; however, detection of ribosome-like structures could provide a robust and sensitive method to seek extant life in combination with other methods.

**One Sentence Summary:** Life, defined as a chemical system capable of Darwinian evolution, likely requires an apparatus to translate heritable instructions into cellular machinery, and we propose to detect this as a biosignature of extant life beyond Earth.

## 1. Introduction

Life is commonly defined as a “self-sustaining chemical system capable of Darwinian evolution” (Benner, 2010; Joyce, 1994; Luisi, 1998; Neveu et al., 2018). In all known forms of life, Darwinian evolution occurs via heritable changes encoded into molecular features (i.e., nucleic acids), which are then translated into cellular machinery and other proteins. These products of translation are responsible for a variety of biological functions within and outside the cell including maintenance, growth, reproduction, regulation, structure, storage, and transport.

In all known life, the ribosome is integral to the “central dogma of biology,” which traces the flow of genetic information into organismal structure and function: genetic information is stored in DNA, which is transcribed into RNA, which is then translated into protein. Specifically, the ribosome is responsible for the process of translation, using mRNA-encoded information to synthesize proteins. All known life is descended from a common ancestor (Pace, 2001), and the structure and genetic code of the ribosome is highly conserved across evolutionary time and the tree of life. Indeed, the most highly-conserved sequences across biological systems correspond to 16S and 23S ribosomal subunits and transfer RNAs, and these sequences have experienced little change over the past 4 billion years (Bray et al., 2018; Isenbarger et al., 2008). Ribosomes are found within all cells regardless of place in the tree of life or level of activity, and are instrumental for a cell’s ability to carry out life-sustaining processes such as growth, reproduction, and metabolism (Bernstein and Baserga, 2004; Bowman et al., 2020).

This conservation of ribosome-associated genomes has been posited as evidence for the existence of an RNA-protein world before DNA became the dominant form of nucleic acid-based information storage (Fournier et al., 2010; Goldman et al., 2010; Harish and Caetano-Anollés, 2012; Hsiao et al., 2009; Jerome et al., 2022). This “RNA world” model for the origin of life posits that RNAs may have acted as both heritable material as well as cellular machinery prior to the evolution of the ribosome or other more sophisticated biological structures. This feature persists today, with RNA molecules being capable of both storing genetic information and catalyzing biochemical reactions. However, there is good reason to assume that life beyond Earth or an alternate origin of life could utilize a heritable storage system that is physically distinct from translation-performing cellular machinery, as discussed in section 2.1.

Translation-performing molecules similar to ribosomes could function as an agnostic biosignature, providing evidence for “life as we don’t know it” that may not share a common origin or physiochemical basis with life on Earth. Given the uncertain range of possibilities for alien biochemistry, morphology, and other characteristics, various methodologies to search for non-terran lifeforms have been proposed that are not specific to known biochemistry. Such strategies can identify potential agnostic biosignatures via direct detection, such as using DNA probes (apatamers) to bind to repeating patterns within compounds (Johnson et al., 2018). They can also be elucidated by analyzing observed molecules: for example, the probability of a molecule of a certain complexity forming abiotically can be quantified (Marshall et al., 2021), and identification of such molecules can be accomplished via mass spectrometry (Chou et al., 2021). Such methods generally rely on the assumption that observation of abundant molecules with sufficiently high complexity can indicate a biotic origin.

Molecules with specific structural or functional features have been proposed as potential agnostic biosignatures. Such features are generally analogous to what is known about terran life without presupposing any origin or biochemical basis. For example, information-encoding charged linear polymers could serve as a mechanism to store, regulate, and transmit genomic information, analogous to nucleic acids on Earth (Benner, 2017). We extend this point, and argue that the “Darwinian evolution” definition of life not only implies a mechanism to store heritable information, but further necessitates a mechanism to translate this information into material that is useful for the biological system. In terran life, this need is satisfied by the ribosome, a molecule that is composed of RNA and proteins and is responsible for using genetic information to synthesize polypeptides and proteins through a process known as translation. The ribosome has a very specific structure and size (although it is important to note that eukaryotic ribosomes are 25-50% larger in diameter), and we similarly argue that non-terran life may use a molecule of similar structure and size. Here, we demonstrate that DNA and RNA can be detected with solid-state nanopore instrumentation. We also describe the case for targeting the translational apparatus as a biosignature, and demonstrate the ability to target our only known example, the ribosome, using solid-state nanopores. Finally, we show that machine learning can discriminate between different classes of biomolecules (DNA, RNA, and ribosomes) based on the features associated with their solid-state nanopore detections.

## 2. The Case for the Translational Apparatus as a Biosignature

### 2.1 What would life look like without translation?

Any life with separate heritable material and cellular machinery must have a translation system to convert instructions into machinery. If these functions are not separate, then heritable material must also act as machinery, including both self-copying and carrying out all other life functions (metabolism, maintenance, growth, evolution). While ribozymes have been discovered and engineered that are capable of copying short RNA sequences (Been and Cech, 1988; Ekland and Bartel, 1996; Horning and Joyce, 2016; Sczepanski and Joyce, 2014), and RNA sequences can be inherited via viral replication (Rampersad and Tennant, 2018), we are not aware of any truly self-replicating informational polymer system of non-trivial complexity (i.e., capable of replicating nucleic acid strands longer than tens of bases). Furthermore, such systems would be unlikely to competitively support the chemical richness required to subsume all the roles of cellular machinery (e.g., an RNA-only RNA world in the absence of proteins). In other words, if such a system evolved, it would likely be displaced by a system with separate coding and machinery due to selective pressures, just as may have happened with life as we know it (Gilbert, 1986; Orgel, 1968).

Separation of coding and machinery would decouple conflicting functions. To perform a wide range of complex functions, cellular machinery requires chemical diversity, whereas enabling copying requires chemical simplicity to facilitate copying robustness. Even while RNA can serve as both as an informational storage molecule and as cellular machinery (as evidenced by ribosomes, tRNAs, and other ribozymes), a physically separate translation apparatus might be expected to arise because decoupling these two functions offers significant benefits, as observed by these functions being decoupled in all known life.

Furthermore, separation among classes of functional biomolecules allows for their maintenance and enables structural differences that align with function. For example, linear charged polymers are considered as a potential universal heritable information storage format for aqueous life (Benner, 2017). In terran life, such polymers exist as nucleic acids, which store genetic information in linear base sequences. Their topology contributes to copying fidelity: branched or otherwise more complicated structures would pose difficulties for effective copying. Additionally, the charge of nucleic acids separates the properties of the molecule from its information content, so that the molecule can be effectively copied regardless of information content. Conversely, the ribosome has a complicated and specific structure that aids in its preservation and enables its highly specialized role in translation. The assembly of two types of biopolymers (polynucleotides and polypeptides) yields a functional and stable ribosome capable of conducting its important catalytic role, leveraging the qualities of both nucleic acid and protein to enable translation (Runnels et al., 2018). The differing charges across the ribosome similarly allows it to function by enabling compatibility with the biomolecules that interact with various parts of the ribosome (Leininger et al., 2021). Furthermore, self-complementarity (the propensity for a biomolecule to bind with itself) among biopolymers enables self-preservation and functional separation of different types of biomolecules. Meanwhile, heterogenous assemblages of different types of biopolymers can beget more complex functionality and “partner-preservation”, where mutualistic assemblies of different biomolecule types can confer protective effects on each other (Runnels et al., 2018). All known life requires a translation apparatus. However, let us consider life that does not have this layer of abstraction, such that hereditary molecules also serve as cellular machinery. As alluded to above, this introduces complexity into copying: changes in information content would result in changes in machinery, which could overly constrain such life’s exploration of evolutionary space. In short, life without translation would be limited in complexity, capabilities, and evolvability (Cuevas-Zuviría et al., 2023), the latter of which is central to our definition of life.

### 2.2 Detection Approaches

A classic approach for detection and identification of bacteria and archaea involves using polymerase chain reaction (PCR) to amplify and sequence the gene encoding for 16S ribosomal RNA (rRNA), which is highly conserved across prokaryotes (Isenbarger et al., 2008). This genomic approach is integral for the study of microbial communities on Earth (Kim and Chun, 2014), since it targets a genomic sequence that exists in nearly all bacteria and archaea, or for some choices of primers, certain large clades of organisms. Similarly, the eukaryotic 18S rRNA gene can be used to identify study eukaryotic community members. Ribosome profiling techniques can be used to sequence mRNA fragments that are being actively translated, enabling study of translation with single-cell resolution (Heiman et al., 2008; VanInsberghe et al., 2021). However, targeting specific ribosome-associated RNA sequences is only suitable for seeking lifeforms that are ancestrally related to known life. One instance where this may be relevant would be for searching for Martian microbial communities that share an origin with life on Earth, e.g., transferred between the planets via lithopanspermia (Carr, 2022; Mileikowsky et al., 2000).

Analysis of rRNA can be used to identify active taxa and assess activity in environmental microbial communities, although caveats exist for these use cases. The relationship between rRNA concentrations and microbial activity is complex and therefore simple correlations do not often exist between the two, and any existing relationships are not consistent across taxa (Blazewicz et al., 2013). Furthermore, in certain environmental conditions (e.g., low temperature, low water activity, high osmolarity), the rRNA itself may persist after environmental conditions have changed, influencing conclusions about which specific microbes are actually currently active in an environment (Schostag et al., 2020). Although drawing explicit conclusions about which microbial taxa are currently active is difficult from rRNA analysis alone, detection of ribosomal material can serve as an indicator that microbial activity has occurred recently or is actively underway in a habitat.

Content-agnostic approaches such as can be used to detect ribosomes without making assumptions about their genetic associations. Microscopy techniques, including fluorescence microscopy and cryoelectron microscopy, can provide information about ribosomal structure and morphology, location and organization within cells, and study of translation (Bai et al., 2013; Bakshi et al., 2012; Fu et al., 2011; Murphy et al., 2020). However, such technologies are destructive, require extensive sample preparation, or are susceptible to image artifacts. Additionally, the size, sensitivity, and complexity of these instruments present difficulties for adapting these types of microscopes for *in situ* space exploration applications.

Dynamic light scattering (DLS) can be used to characterize molecules in an aqueous sample by measuring their diffusion coefficient, which in turn can be used to infer molecule size and shape (Stetefeld et al., 2016). For example, DLS can be used to interrogate ribosomal shape (Bruining and Fijnaut, 1979), track structural changes (Patel and Cunningham, 2002), and make comparisons between ribosomal types (Müller et al., 1986). DLS is a good strategy for making accurate size and shape particle measurements in minutes with a small amount of sample (<3µL). Additionally, sample preparation is minimal, and this technique is compatible with high-concentration and turbid samples. However, measurements are sensitive to temperature and solvent viscosity, and DLS is not capable of achieving sufficiently high resolution to distinguish between closely-related molecules, such as monomers vs. dimers. Finally, DLS is sensitive to changes in size and concentration (Jose et al., 2019). Overall, DLS is a good technique for interrogating pure ribosomal samples at high concentration, but is not viable for identifying or characterizing ribosomes within a heterogenous sample or when present at low abundance.

Solid-state nanopores (SSN) can provide a useful method for ribosomal detection with single-molecule or single-particle resolution. This instrumentation detects individual molecules or particles (hereafter referred to in the generic sense as particles) in an aqueous sample of known conductivity by applying an ionic current to a nanometer-scale pore in a synthetic membrane, using a voltage or pressure differential to pass particles through the pore, then measuring how the particles disrupt the ionic current. The magnitude of the change in conductance and duration of the blockage can then be used to determine the relative size of each particle (**Figure 1**). SSN instrumentation is distinct from biological nanopore sequencing (e.g., the MinION from Oxford Nanopore Technologies), which uses controlled translocation of nucleic acids regulated by biological components with specific base sequences inferred from changes in ionic current flow. Instead, due to its larger, non-specific pore, SSN tools can analyze a wide size range of particles and are not constrained to a specific class of molecules. This benefit is also a challenge to achieving molecular specificity and to achieving single base resolution, as discussed below.

**Figure 1.**
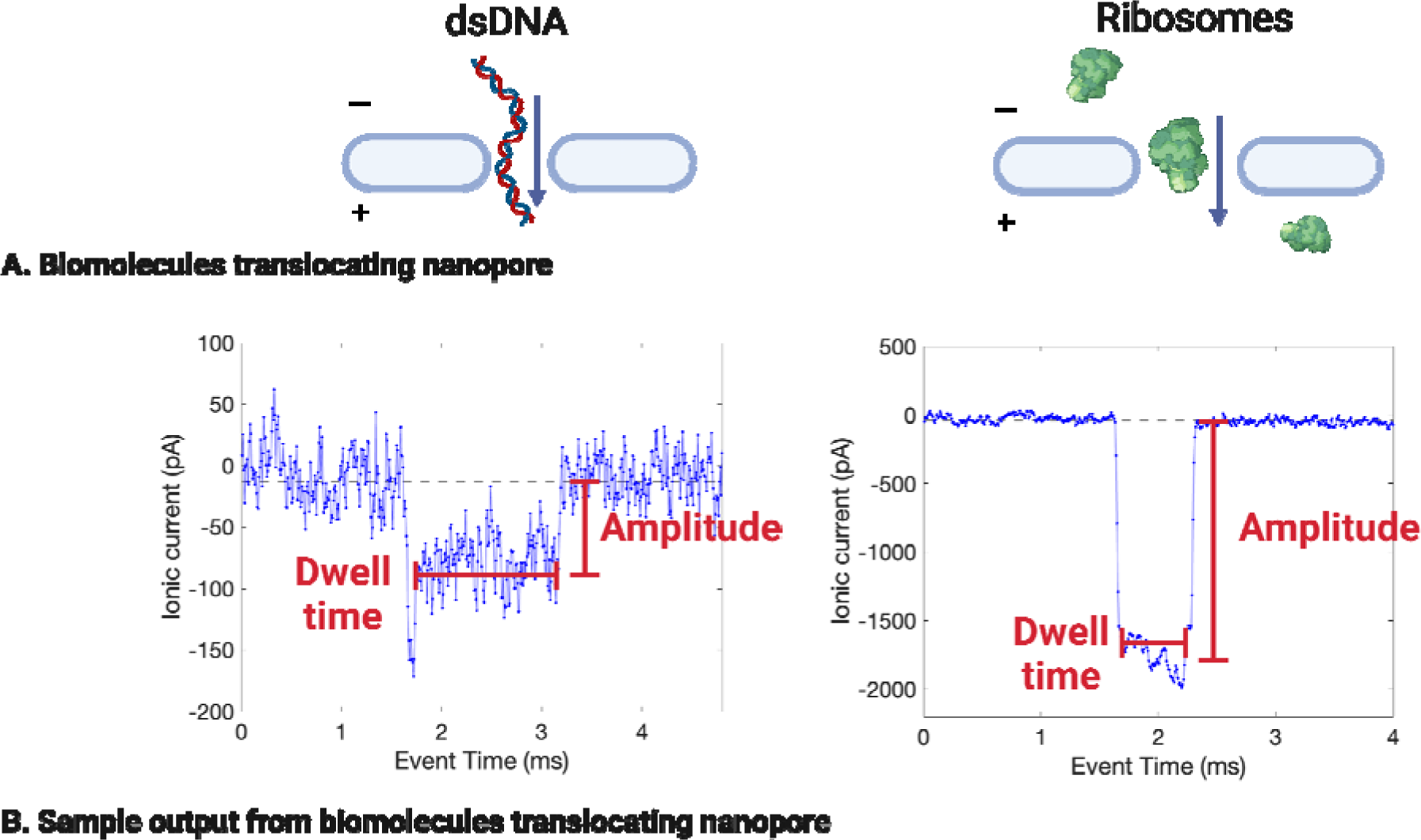
Examples of DNA and ribosome detection using a solid-state nanopore (SSN) in 2M LiCl. Molecules passing through a SSN (A) cause momentary blockages in the membrane’s ionic current (B). In later analysis, the change in amplitude is reported as the change in conductance (since the voltage applied is constant). Note the order of magnitude difference in the vertical axis.

In recent years, major advancements have been made in SSN technologies to improve their sensitivity, selectivity, and throughput capacity, enabling development of applications in biology, chemistry, and medicine (Xue et al., 2020). Particularly of interest to this work and other astrobiological applications, SSN-based methodologies have been used to detect biomolecules such as nucleic acids and ribosomes in laboratory conditions. Raveendran et al. (2020) used nanopipettes fitted with ∼60 nm nanopores on the tips to detect eukaryotic 80S ribosomes (derived from human and *Drosophila melanogaster* cell cultures) to detect ribosomes and distinguish ribosomes from polysomes (groups of ribosomes bound together by mRNA strands). Rudenko et al. (2011) used a nanopore-microfluidic chip to detect individual prokaryotic 50S *Escherichia coli* ribosomes and control their rate of nanopore translocation. Similarly, Rahman et al. (2019) used an optofluidic chip to control the flow of ribosomes, DNA, and proteins through a nanopore and measure these biomolecules as they translocated the nanopore. Xia et al. (2022) used a nanopore drilled into a silicon nitride membrane on a glass chip to detect and characterize DNA that was extracted from Martian analog soils, directly demonstrating SSN applications for astrobiological life detection.

Machine learning (ML) can be applied to solid-state pore measurements to classify and identify biological particles, given appropriate training data. Xia et al. (2021) applied multiple ML models to their SSN data sets to differentiate signals of monosaccharides and disaccharides, which enabled them to determine the composition of polysaccharide compounds. Arima et al. (2020) achieved over 99% accurate classification of viruses using ML. Using a molybdenum disulfide membrane, instead of the commonly-used silicon nitride membrane, Farimani et al. used ML to identify individual amino acids within polypeptide chains (Farimani et al., 2018). On the micropore scale, ML can be similarly used to identify and classify cells. Using solid-state micropore data, Hattori et al. applied ML to discern between five clinically-relevant bacterial species based on physical features including cell shape, surface charge density, mass, surface proteins, and motility (Hattori et al., 2020). Tsutsui et al. tagged bacterial walls with synthetic peptides and analyzed them with a solid-state micropore, then used machine learning to identify bacterial features and discriminate between strains (Tsutsui et al., 2018).

Solid-state nanopores are non-specific and can be used to detect not just biomolecules such as nucleic acids and ribosomes, but also non-biological particles. Due to this non-specificity, it has been proposed that solid-state nanopores may be useful for detecting agnostic biosignatures (Bywaters et al., 2017). It has also been proposed that alien lifeforms with a separate origin and alternative biochemistry from terran life may store genetic information in charged linear polymers that are structurally and functionally analogous, but not necessarily chemically similar, to nucleic acids (Benner, 2017). We argue here that translational machinery analogous to ribosomes may be present and could additionally serve as an agnostic biosignature. Solid-state nanopore instrumentation can detect and characterize both nucleic acids and ribosomes, and hold potential for detecting similar structures produced through alternate biochemistries.

Here we explore the utility of ribosomes (and agnostic translational machinery) as a biosignature, and demonstrate nucleic acid and ribosomal detection using solid-state nanopore instrumentation. Additionally, we propose that detecting one or more peaks indicative of translational machinery, in a sample that has been agitated to sufficiently lyse any existing cells, could indicate the presence of life.

### 2.3 Expected Size of Translational Apparatus for Life as We Don’t Know It

Life beyond Earth, even if ancestrally or chemically distinct from life as we know it may utilize a structure similar in size to the ribosome, balancing the energetic costs of assembling translation machinery with the minimum complexity required for such a machine. In short, the translation machinery must have some minimum size related to the complexity of its translation function. In addition, there is likely a selective advantage to the translation machinery not being far larger than the minimum size, due to the energetic and time cost of synthesis and assembly (Li et al., 2014; Shore and Albert, 2022). Additionally, different life forms within the descendants of a particular origin of life may have translation apparatus that differ in size, similar to how prokaryotic and eukaryotic ribosomes have different sizes on Earth. The ancient ribosomal core, which was capable of translation and has been evolutionarily conserved since before the last universal common ancestor (LUCA), is about 2 MDa in size. Contemporary prokaryotic ribosomes are about 2.5 MDa, with a slightly larger size due to additional proteins and a slightly longer rRNA sequence (about 100 nucleotides larger than the core) (Bowman et al., 2020). Smaller bacterial ribosomes also exist, with certain ribosomal proteins being absent in bacteria with exceptionally small genomes (Nikolaeva et al., 2021). Eukaryotic ribosomes range in size from about 3.5 to 4 MDa, similarly attributable to longer rRNA sequences (hundreds to thousands of nucleotides larger than the core) and additional proteins (Bowman et al., 2020). It has been proposed that the size difference between prokaryotes and eukaryotes reflects additional complexity in eukaryotic regulation of translation (Lafontaine and Tollervey, 2001). Comparisons of eukaryotic and prokaryotic ribosomes can allow for identification of the common ancestral core, and molecular models can be used to infer how ribosomes may have evolved (Petrov et al., 2014). Modeling suggests that various ribosomal capabilities for interpreting genetic encoding and creating polypeptides in response were sequentially added, resulting in a molecule composed of functional proteins and nucleic acids (Petrov et al., 2015). The tight size distribution and number of ribosomal proteins (∼50 in prokaryotes, ∼80 in eukaryotes) may reflect selection for optimal duplication efficiency (Shore and Albert, 2022). We do not presume any chemical specificity for ribosome-like machinery; rather, we argue that function and approximate size may be conserved across multiple origins of life with molecules that may differ in structure or composition.

## 3. Methods

### 3.1 Reagents and Reagent Preparation

Aliquots of thawed biomolecule samples (single-stranded RNA [ssRNA] ladder [NEB #N0362], 1 kilobase [kb] double-stranded RNA [dsRNA] ladder [NEB #N0363], 1 kilobase double-stranded DNA [dsDNA] ladder [Thermo Scientific #SM0311], high range dsDNA ladder [Thermo Scientific # SM1351], DNA plasmid pUC19 [NEB #N3041], *E. coli* ribosome [NEB P0763S]) were added to Ontera Start-Up Buffer (2M LiCl, conductivity 11.96 S/m).

### 3.2 Solid-State Nanopore Operation

The Ontera NanoCounter was operated using the associated software (NanoCounter v0.19.1). Nanopores were conditioned following manufacturer instructions, using the Ontera Start-Up Buffer (2M LiCl, conductivity 11.96 S/m) as the conditioning buffer. Biomolecule samples and negative controls were loaded into nanopore and analyzed following manufacturer instructions.

### 3.3 Detection of dsDNA, ssRNA, dsRNA, Ribosomes

For a given experiment, the same pore was loaded with a sequence of samples after conditioning, using an alternating pattern of negative control (0.2µm-filtered buffers without added biomolecules) (run time 3-5 minutes), followed by a sample (run time 6-10 minutes), followed by the next negative control. This was done to assess buffer cleanliness, flush out and assess any carryover from prior samples, and quantify the risk of false positives. The NanoCounter software used an automatic event extraction program to identify events with a signal-to-noise ratio high enough to consider the event to be a particle translocating the nanopore.

### 3.4 Application of Machine Learning to Differentiate Biomolecule Types

To prepare the SSN data for machine learning classifier training, event metadata was compiled for all biomolecule events with a signal-to-noise ratio (SNR) greater than 5. The MATLAB Classification Learner app (with MATLAB_R2023a) was used to train several machine learning algorithms on the data. Data was partitioned by the toolbox, with 25% used for holdout validation, 10% used for testing, and the remining 65% used for training. The response variable was the sample identifier (“sample_name”), and the predictor variables were dwell time (“dwell_sec”), signal-to-noise ratio (“SNR”), mean current amplitude during event (“mean_amp_pA”), maximum current amplitude during event (“max_amp_pA”), median current amplitude during event (“med_amp_pA”), standard deviation of amplitude (“std_amp_pA”), and area of the event disruption (“area_pA_sec”). The Ontera Nanocounter software recorded current values in picoamps and associated conductivity values in nanosiemens; since the applied voltage was constant across experiments and thus conductivity always scaled with current, only current values were used for the machine learning training to avoid redundancy.

## 4. Results

### 4.1 Solid-State Nanopore Detection Achieved a Low False Positive Rate

Negative controls were run before every test sample to assess background and carryover from previous runs. When testing sterilized and filtered brines on new nanopores, event abundance was low (<10 events per minute). Events detected in this case are likely particle contaminants introduced during buffer preparation, sample loading, or present within the flow channels of the nanopore. Event abundance in the negative controls when ran between DNA samples was similarly low, as shown in **Figure 2**. A higher abundance of events than usual was sometimes detected in pores when running negative controls between biomolecule samples (**Table S1**). This was especially prominent in cases where clogging (for example, high-concentration ribosome samples) and biomolecule precipitation (for example, RNA in lithium chloride) were suspected to occur.

**Figure 2.**
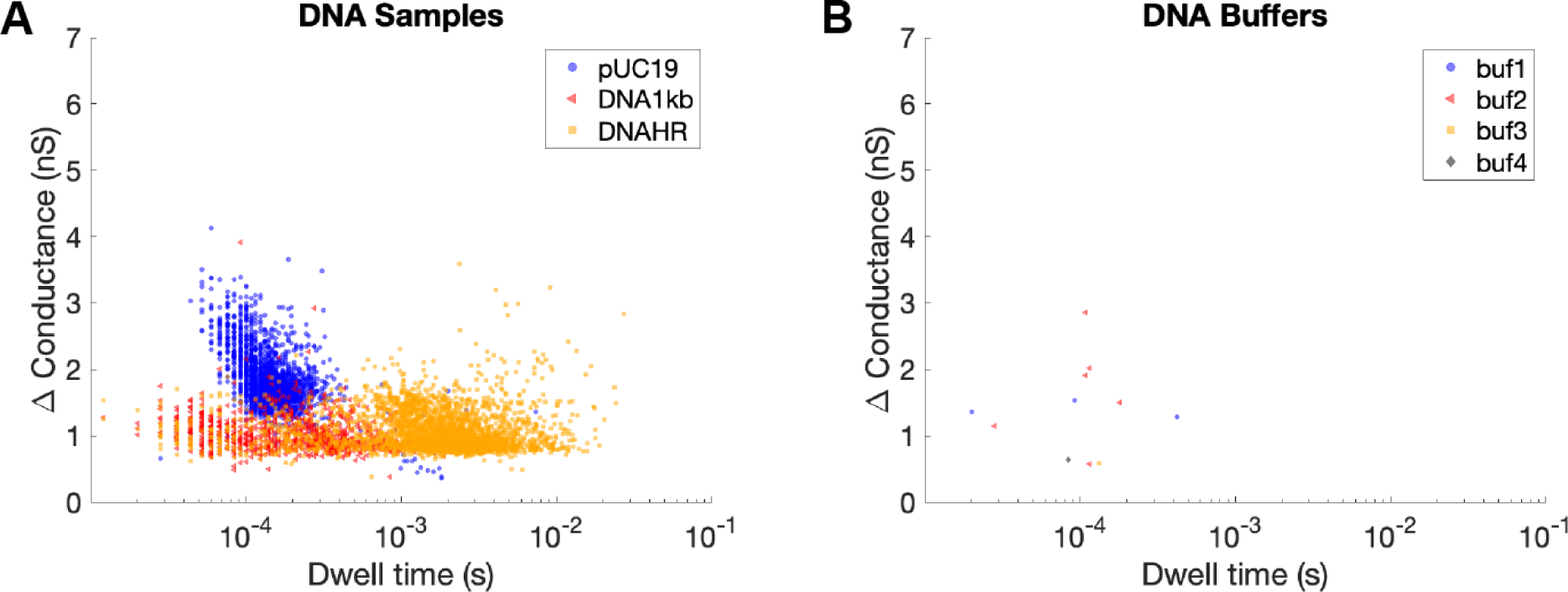
Overlaid event data (conductance vs. dwell time) for biomolecule (A) and buffer (B) samples in 2M LiCl. Buffer and biomolecule samples were run in the following order: buf1, pUC19, buf2, DNA1kb, buf3, DNAHR, buf4 (**Table S1**). Buffer (negative control) event counts demonstrate low carry-over between different samples as evidenced by low (<10) event abundance for run times of approximately 3 minutes for each buffer, and 6-10 minutes for each biomolecule sample.

### 4.2 Different Biomolecule Classes and Sizes Have Distinct Signatures

DNA and RNA samples were suspended in 2M LiCl and detected using the Ontera NanoCounter. The DNA samples were plasmid pUC19 (a circular fragment with a length of 2,676 base pairs), 1 kb dsDNA ladder (linear DNA fragments of 14 different sizes ranging from 250 to 10,000 base pairs), and high range dsDNA ladder (linear DNA fragments of 8 different sizes ranging from 10,171 to 48,502 base pairs). The RNA samples were single-stranded RNA (ssRNA) ladder (linear ssRNA fragments of 7 different sizes ranging from 500 to 9,000 bases) and double-stranded RNA (dsRNA) ladder (linear dsRNA fragments of 7 different sizes ranging from 21 to 500 base pairs). Events were automatically extracted from these runs using the NanoCounter software, totaling 2,284 events over 6.62 minutes for plasmid pUC19, 919 events over 5.93 minutes for 1 kb dsDNA ladder, 3,115 events over 5.99 minutes for high range dsDNA ladder, 1,874 events over 10.03 minutes for ssRNA, and 1,398 events over 11.01 minutes for dsRNA (**Table S1**). Key features were also identified, most importantly dwell time (the amount of time that the molecule spent translocating the nanopore) and change in conductance (resulting from the current disruption of the molecule translocating the nanopore). Given the low background noise when running biomolecule-free buffer solutions (**Figure 2, Table S1**), we assume that all events from these runs are indeed the target biomolecule translocating the nanopore.

DNA events displayed specific features that were indicative of biomolecule size and structure (**Figure 3**). The change in conductance for pUC19 was generally approximately double that of the HR and 1kb ladders. This is thought to be attributed to 4 strands (2 dsDNA strands) passing through the nanopore at once for a circular DNA plasmid, as compared to the 2 strands (1 dsDNA strand) that would pass through at once with a linear ladder. The lower-range ladder (1kb) had a shorter range of dwell times compared to the upper-range ladder (HR), which is indicative of the respective ladder fragment lengths.

**Figure 3.**
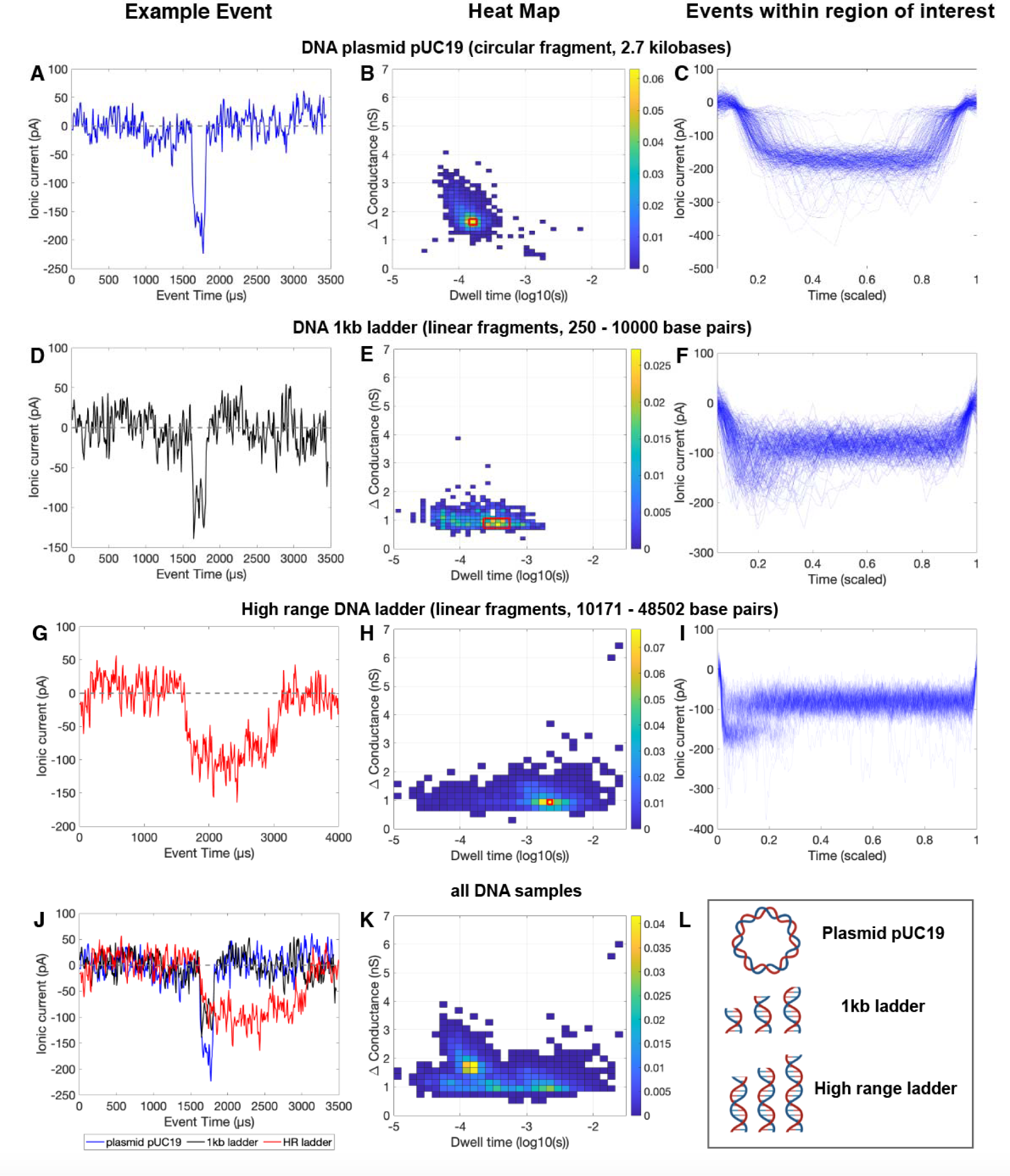
Event data for DNA plasmid pUC19 (A-C), 1 kilobase ladder (D-F), high-range ladder (G-I), and all DNA types (J-K) in 2M LiCl. Left panels (A, D, G, J) show selected event examples. Middle panels (B, E, H, K) show heat maps of change in conductance vs. dwell time (time molecule spends disrupting nanopore’s ionic current). Heat map color scale correspond to probability of an event being in that region of the heat map. Red boxes indicate regions of interest for right panels. Right panels (C, F, I) show all overlaid events within red boxes with a signal-to-noise ratio greater than 5, with time scaled to 1. Panel L shows schematic representations of the three measured types of DNA. In Panel K, each probability distribution (pUC19, 1kb ladder, high range ladder) is evenly weighted to facilitate comparison between the different classes.

The ssRNA and dsRNA events also differed in their change in conductance and dwell time, with ssRNA samples generally having a slightly larger change in conductance and slightly smaller dwell time than the dsRNA samples (**Figure 4**). However, these differences are less discernable for the RNA samples than the DNA samples. This may be attributable to the differences in molecule size and structure between the two types of DNA and RNA tested: the two DNA samples differed in conformation (circular plasmid of a specific length vs. linear fragments of varying lengths), while both RNA samples consisted of linear fragments spanning similar ranges of sizes but in double or single stranded form. Strand number or topology can be expected to influence persistence length and thus secondary structure (e.g., ssRNAs are highly flexible). Additionally, high numbers of events in the buffers immediately after running RNA suggests that precipitation and resolubilization may have occurred for these samples, which could similarly result in low-quality events due to interactions with the nanopore.

**Figure 4.**
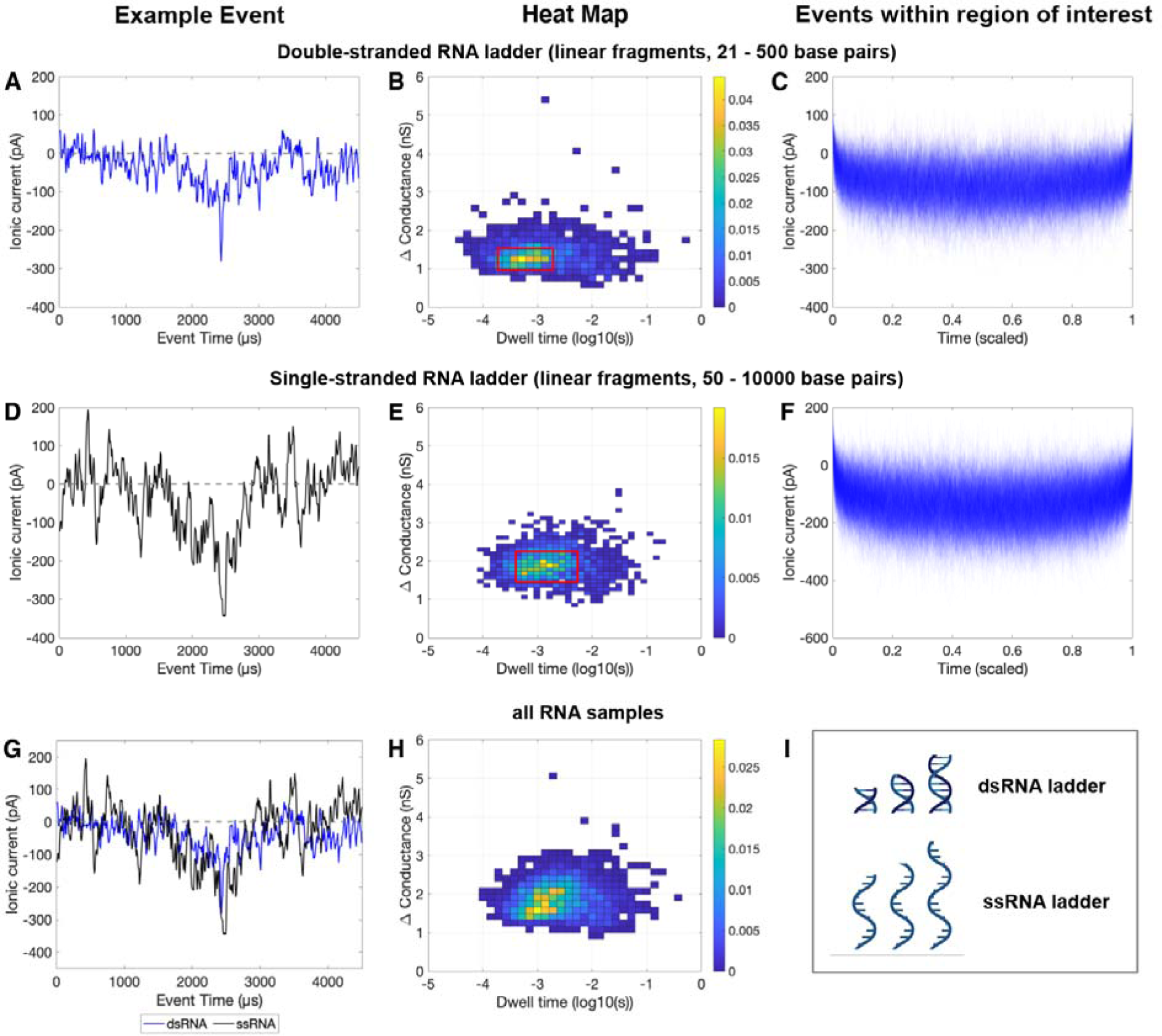
Event data for RNA samples: double-stranded RNA (A-C), single-stranded RNA (D-F), and both RNA types (G-H). Left panels (A, D, G) show representative events. Middle panels (B, E, H) show heat maps of change in conductance vs. dwell time (time molecule spends disrupting nanopore’s ionic current). Heat map color scales correspond to probability of an event being in that region of the heat map. Red boxes indicate regions of interest for right panels. Right panels (C, F) show all overlaid events within red boxes, with time scaled to 1. Panel I shows schematic representations of the two measured types of RNA. In Panel H, each probability distribution (dsRNA ladder, ssRNA ladder) is evenly weighted to facilitate comparison between the different classes.

Similarly, *E. coli* ribosomes were suspended in 2M LiCl and measured with the Ontera NanoCounter. 17,077 events were extracted from 10.03 minutes of run time (**Table S1**). **Figure 5** contains a heat map of events from this run (Panel A), individual examples of events interpreted to be an intact ribosome and a ribosomal fragment (Panel B), and overlaid events from the regions of the heat map interpreted to be intact ribosomes (Panel C) and ribosomal fragments (Panel D). Of the 17,077 events, 6,885 are thought to represent intact ribosomes translocating the nanopore (defined as events with a change in conductance greater than 8 nS). The large number of events with changes in conductance smaller than 5 nS are thought to be ribosomal fragments. Within this results space, there is a high concentration of events with a change in conductance between 1-5 nS and a dwell time between 0.00001-0.0001 seconds. These are interpreted as protein fragments. Some events have a larger dwell time (0.0001-0.1 seconds), which closely aligns with the reign occupied by RNA events (**Figure 4**). Thus, it is thought that these fragments may be RNA products of ribosome fragmentation.

**Figure 5.**
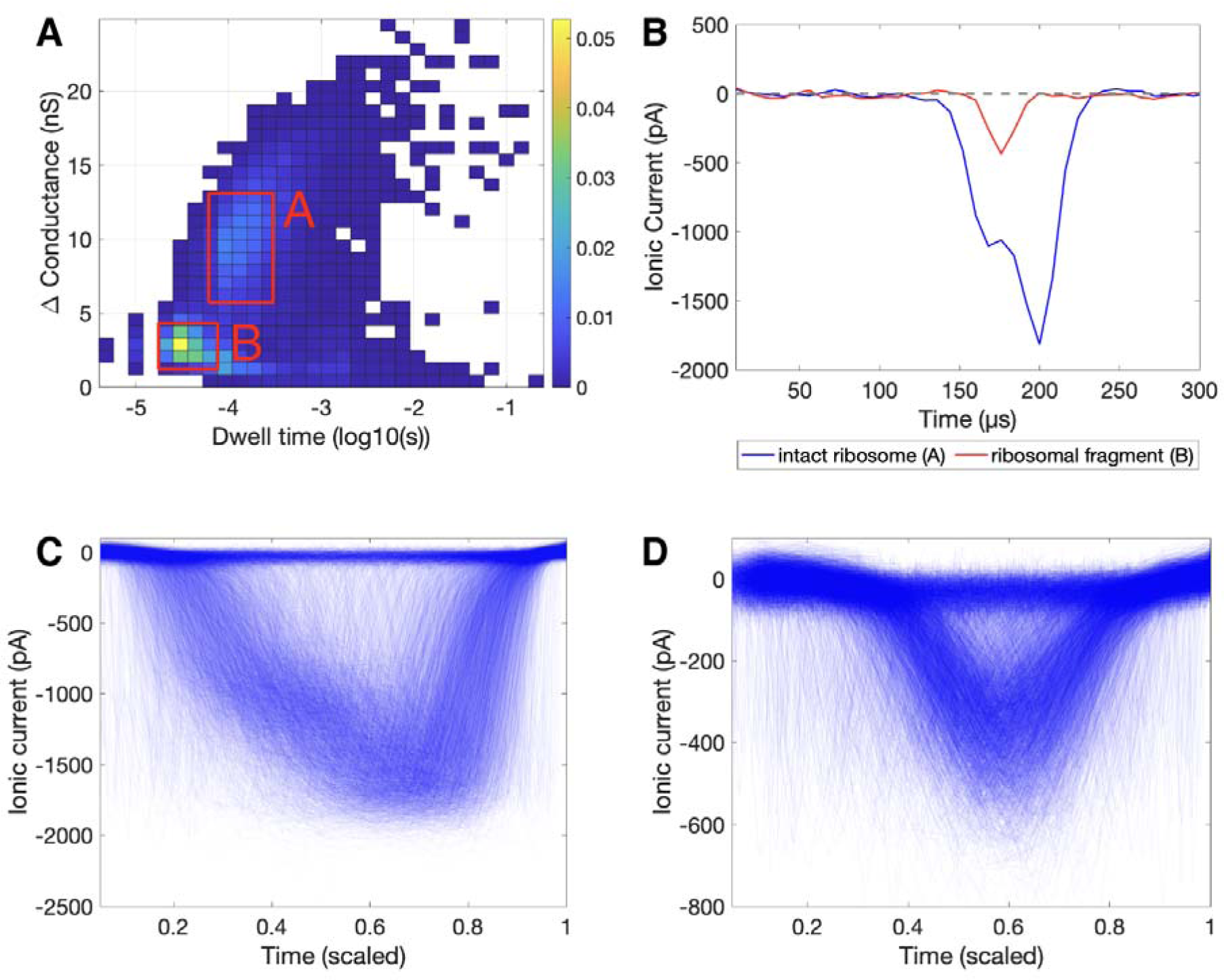
Event data for *E. coli* ribosome in 2M LiCl. Panel A shows heat map of change in conductance vs. dwell time (time molecule spends disrupting nanopore’s ionic current). Heat map color scale corresponds to a probability of an event being in that region of the heat map. Panel B shows example events interpreted to be an intact ribosome (corresponding to Panel C) and a ribosomal fragment (corresponding to Panel D). Panels C and D show overlaid events within the red boxes C and D on Panel A, plotting all events with a signal-to-noise ratio greater than 10, with time scaled to 1. Event data around a current baseline of 0 pA can be observed in C and D throughout the middle of the scaled events (e.g. events with a current signal close to the baseline along time axis points of 0.4 - 0.6). This is due to instances of the automatic event extraction selecting an event with 2 translocations (due to the translocations happening close to each other temporally).

Overall, based on dwell time and change in conductance, signals from each biomolecule class cluster together, although there is some overlap (**Figure 6**). Machine learning was implemented in attempts to better differentiate between sample types.

**Figure 6.**
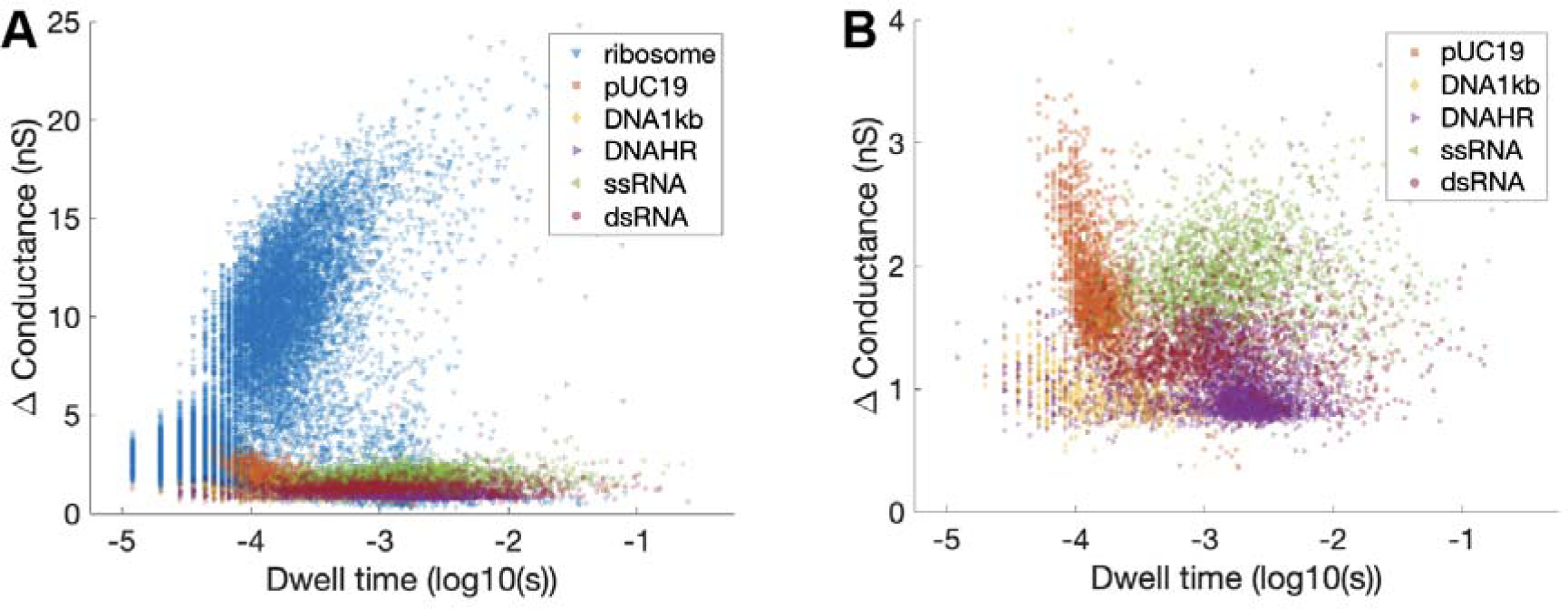
Scatter plots of conductance vs. dwell time for various biomolecule types, sizes, and conformations. Panel A includes all ribosome, DNA, and RNA samples. Panel B exclude ribosomes so that distribution of DNA and RNA events can be seen more clearly.

### 4.3 Machine Learning Algorithms Can Differentiate Biomolecule Types and Structures

The Classification Learner app identified the validation and testing accuracy of a variety of machine learning classifier methods, including neural networks, support vector machines, k-nearest neighbors, kernel approximation, naïve Bayes, discriminant, logistic regression, and tree-based models (**Table S2**). To provide an illustrative example, detailed results from the medium tree model are shown. This specific model was selected for its high interpretability, as other top-scoring algorithms (e.g. neural networks, support vector machines, k-nearest neighbors) can be more difficult to interpret. The medium tree was specifically selected from the applied decision tree models (fine tree, medium tree, coarse tree) as it balances ease of interpretation with accuracy – the fine tree model had higher accuracy levels (94.41% validation accuracy and 95.02% testing accuracy, compared to the medium tree’s 92.89% validation accuracy and 93.34% testing accuracy), but the fine tree involved a much more complex array of decisions. Additionally, with limited training data, it is difficult to preclude overfitting, which is more likely to occur for finer trees. The medium tree (**Figure S1**) contained 19 splits to classify biomolecules based on event features (**Table S3**). It is possible to reach the same biomolecule classification by following different paths throughout the tree, which may be attributable to the presence of different molecule morphologies within a single sample (e.g. intact vs. fragmented ribosome, supercoiled vs. unsupercoiled plasmid, bent vs. straight DNA/RNA fragment). The medium tree’s confusion matrix (**Figure 7**) demonstrates validation performance, showing how biomolecules were generally classified correctly by this model. This specific model is shown here due to its interpretability, but other models achieved higher performance and warrant further exploration.

**Figure 7.**
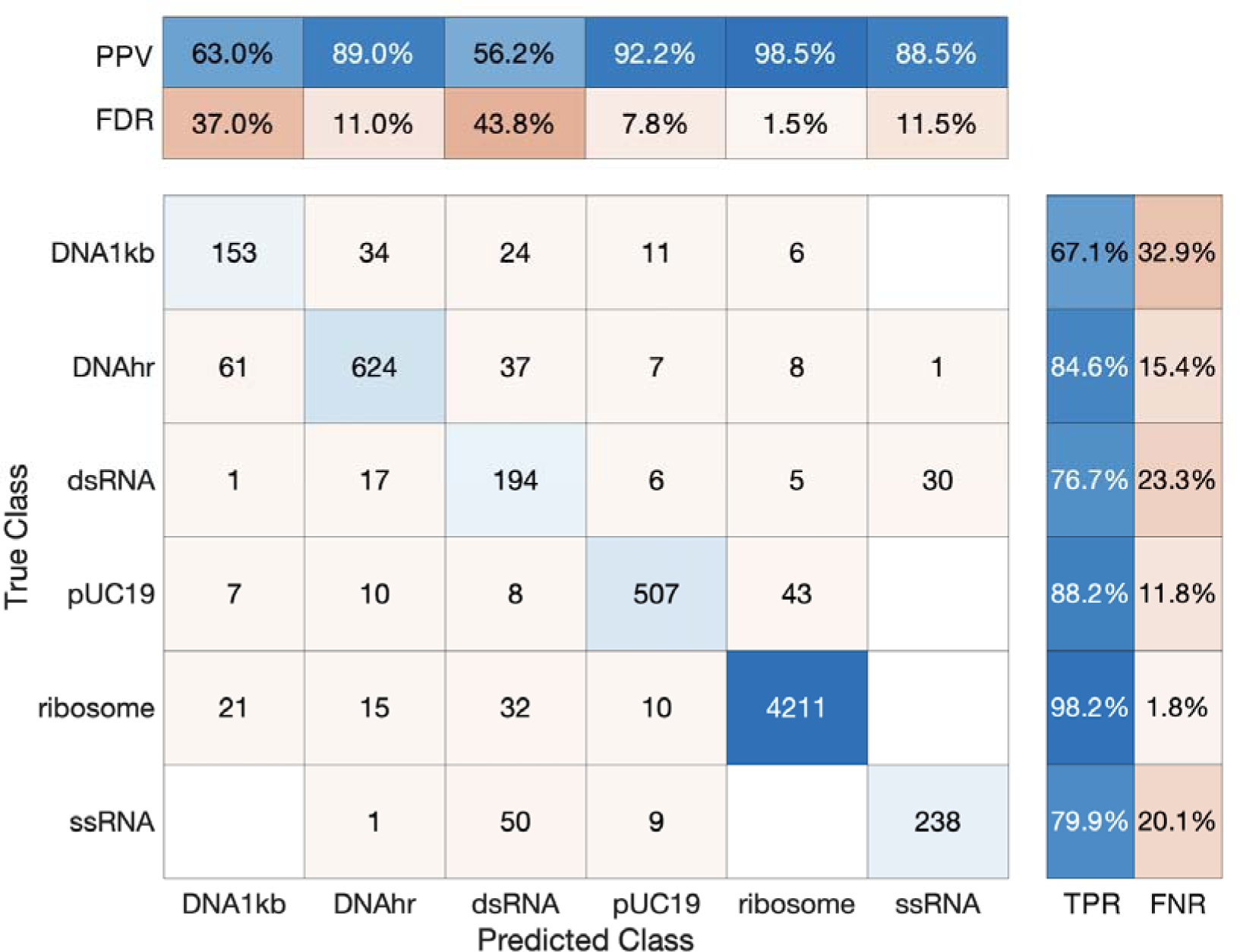
Confusion matrix for Medium Tree model. This matrix was generated by th Medium Tree model by the MATLAB Classification Learner app (MATLAB_R2023a). True Class vs. Predicted Class matrix shows the raw numbers of events for each true/predicted class combination. The upper matrix shows the likelihood that an event is correctly classified given the true class by showing the Positive Predictive Value (PPV), the percentage of correctly classified events within all events of that class, and the False Discovery Rate (FDR), the percentage of incorrectly classified events within all events of that class. The right matrix shows how many of the events were correctly predicted within a predicted class by showing the True Positive Rate (TPR), the percentage of correctly classified events within all events assigned to that class, and the False Negative Rate (FNR), the percentage of incorrectly classified events within all events assigned to that class.

## 5. Discussion

Here, we argue that ribosomes can serve as a biosignature indicative of terran-like life, and another instance of cellular translational machinery of similar size and structure could serve as an agnostic biosignature. In a case of “life as we don’t know it,” a higher abundance of particles within a certain size range compared to the rest of the sample could serve as an agnostic biosignature. Using solid-state nanopore instrumentation, we demonstrate that such cellular machinery can be detected and distinguished from other biomolecules in aqueous samples.

Such measurement is sensitive (capable of correctly identifying target molecules in the sample) as evidenced by successful detection of various biomolecules, and specific (able to discern whether a sample does not contain target molecules) as evidenced by the low abundance of events within the negative controls. When measuring negative controls, no special precautions were taken (e.g. filtration of buffers), suggesting the potential for lowering the false-positive event rate further. Target biomolecules were evident within heat map distributions as specific peaks corresponding to different molecule sizes and conformations. This is especially true for ribosomes, which exist in very specific quantized sizes for prokaryotes and eukaryotes and are often highly abundant, e.g., 25,000 to 31,000 per µm^3^ for active *E. coli* cells, as reported by Bakshi et al. 2012. High abundance of ribosomes is not limited to active cells. Slow-growing ultra-small archaea of volume 0.03 µm^6^ (effective spherical diameter of 0.4 µm) have an average of 92 ribosomes/cell (Comolli et al., 2009), or 3000 per µm^3^. Bacterial endospores contain ribosomes at a similar abundance to active cells, even when mRNA has been degraded (Chambon et al., 1968). This high abundance of ribosomes across different types and activity levels of cells may allow for the distinguishing of ribosomes within cellular lysate or environmental samples treated to lyse cells. Conversely, abiotic material that was similarly treated would be expected to have a more even particle size distribution.

Certain features could confound a search for translational machinery as a biosignature, including other cellular components and abiotic features. The severity and implications of such mistaken identities vary. If another cellular component (e.g., genetic information storage molecules, functional proteins, other cellular organelles) is mistaken as a translational apparatus, then life has still been detected. However, if a particle of abiotic origin is mistaken for a ribosome-like molecule, this could cause a false-positive life detection result. For example, in Earth’s atmosphere, mechanically generated particles tend to be in the coarse-mode (>1 µm) size range, with particle sizes in the nanometer to tens of nanometer range forming from nucleation and thus likely to exhibit a continuous size range rather than high abundance at a specific size. Accumulation of nucleated particles can result in larger sizes (typically hundreds of nanometers). However, in astrobiological contexts, additional caution is warranted. While 99% of dust in the Enceladus plume is expected to be below 10 nm with a peak at 2 nm (Meier et al., 2014), the presence of hydrothermal activities at Enceladus was subsequently inferred from Cassini-detected silica nanoparticles in the 4-16 nm size range (Hsu et al., 2015). One mitigation in this case would be to filter a sample to retain cells and reject free silica nanoparticles prior to lysis. An additional approach would be to heat-treat a sample and observe if the particles are heat tolerant or exhibit denaturation or degradation. Further and varied analysis of any detected molecules would be necessary to discern their composition, function, and potential biogenicity. While outside the focus of this paper, future work should evaluate the potential abiotic production of particles in different settings to assess the risk of false positives.

The results presented here are intended to serve as a proof-of-concept study that lays the foundation for more extensive and robust work, with multiple avenues identified for further experimentation. One factor that warrants further study is buffer-biomolecule compatibility. In this study, it was particularly difficult to achieve low event abundance in the negative control buffer samples ran between RNA and ribosome samples. This is likely due to effects of the lithium chloride buffer on these biomolecules, which is thought to have caused molecular precipitation that hindered flow cell flushing and subsequent molecule detection. Similarly, it was difficult to discern ssRNA and dsRNA events by looking at current disruptions, possibly to due to precipitation of the RNA fragments and/or interaction with the nanopore. It seems that most of the detected events were attributable to true translocations of RNA strands through the nanopore, since the event rate was much higher for the RNA samples than the blank runs immediately before and after (**Table S1**). However, many of these events had a relatively low signal-to-noise ratio compared to what was observed for other samples. Refining buffer and pH compatibility with RNA (and other target biomolecules) is a future target for optimization.

Pore size constrained the size of biomolecules that can be analyzed with this system, and may have played a role in the ribosomal fragmentation observed. This study used nanopores embedded within pre-fabricated flow cells that arrived in a desiccated state, which we then hydrated and subjected to an electrochemical conditioning process that widened and stabilized the pore. This conditioning process would result in a pore with a diameter between 20-45 nanometers, which could not be significantly altered. This was sufficient to allow the passage of single strands of nucleic acids (∼2 nm diameter) and prokaryotic ribosomes (21 nm diameter). It is important to match pore and biomolecule size as closely as possible: if the nanopore is too small, it inhibits free flow of molecules through the nanopore. Conversely, if the nanopore is too large, signals from translocating molecules may be obscured by noise – the closer the size of the biomolecule is to the size of the nanopore (without being large enough to clog the nanopore), the higher the signal-to-noise ratio (Xia et al., 2022). With the nanopores used in this study, we were able to detect nucleic acids, but may be able to achieve stronger signals with a smaller nanopore. Additionally, we were able to detect ribosomes, but also observed the presence of smaller particles thought to be ribosomal fragments. Such fragmentation is hypothesized to have been caused by buffer chemistry and exacerbated by RNA precipitation and clogging of the nanopore. Using a larger nanopore in future studies may help alleviate this issue, especially if coupled with a buffer that is more compatible with ribosomes. Overall, having access to a wide variety of nanopore sizes will be useful for future work with molecules of various sizes, structures, and biochemistry. Future analysis of biomolecules with SSN should also include a variety of saline solutions. Nucleic acids and ribosomes are both structurally sensitive to ionic concentration and composition (Shukla and Mikkola, 2020; Spitnik-Elson and Atsmon, 1969; Tian et al., 2023), and biomolecule translocation itself can also be drastically influenced by ionic conditions (Bell et al., 2016; Kowalczyk et al., 2011). More extensive study of SSN performance under various saline conditions will be necessary to achieve better understanding of how solution composition influences results.

Furthermore, we demonstrate that different biomolecule classes and structures can be differentiated and predicted by decision tree machine learning algorithms with over 90% accuracy on both validation and test data. Both a fine and a medium tree were generated, with the medium tree giving a more comprehensible assessment of how biomolecules could be discerned at the cost of less decisions and lower accuracy, and the fine tree yielding higher accuracy classification by making more decisions, at the cost of a very lengthy list of nodes and the potential for overfitting. Other less-interpretable machine learning models (for example, neural networks) yielded even higher accuracy, further indicating the potential that machine learning classification holds for SSN data. This is useful for summarizing and transmitting data during future missions with highly constrained data budgets, such as at or within Ocean Worlds.

These machine learning results are intended to justify more rigorous future application of machine learning-based classification of solid-state nanopore data, including exploring the utility of various algorithms, hyperparameter tuning, and feature engineering. In addition to this computational work, it will be critical to test these algorithms on datasets from different SSN runs under various conditions and instrumentation to ensure that accurate results are consistently obtained. Since the dataset for each biomolecule class was generated by a single experimental run of that specific class, the experimental conditions (e.g., pore size, buffer chemistry, etc.) may influence event data independently of the influence of the specific biomolecule type. Generation of more data by analyzing different biomolecule types, structures, and sizes under various buffer conditions and with different nanopores will be instrumental in refining such a machine learning-based classification.

There are a few major caveats for applying this laboratory-based detection of terran biomolecules towards astrobiological life detection. First, the utility of ribosomes or ribosome-like translational machinery as a biosignature relies on the assumption that such “life as we don’t know it” synthesizes biomolecules for cellular functioning via translation or an analogous mechanism. Although it is not known how life may exist without ribosomes or a ribosome-like particles, we acknowledge the possibility that alternate biochemistries, cellular structures, and life strategies could be associated with life forms that would not be detectable by searching for translational machinery. Additionally, this work assumes that such a translational apparatus will remain stable during cellular lysis, extraction, and analysis conditions. If translational machinery from an alien lifeform were to be destroyed or otherwise disrupted by such processes, this would increase the chance of false negative results. Finally, unknown abiotic conditions could produce confounding structures with similar signatures, resulting in false positive readings. This possibility highlights the need for stringently using negative controls.

We argue that ribosomal detection could serve as one line of evidence for life, and such measurement could be conducted in conjunction with other life detection analysis. Such detection could be particularly useful as a pre-screening tool for samples, subjecting it to this biologically-agnostic particle measurement technique to determine size distribution and potential biogenicity of samples. This could aid in sample selection or prioritization before passing it forward to more targeted analyses (e.g., DNA sequencing, microscopy, mass spectroscopy, or other compositional analyses). Furthermore, we demonstrate how machine learning can be utilized to classify biomolecules based on various features of translocation events measured by a solid-state nanopore device. Overall, SSN presents one strategy to interrogate various components of a sample and search for patterns or features that may indicate biogenicity. Negative results from an attempt to detect ribosomes would not necessarily preclude the possibility for life forms to be present within the sample. Opportunities for *in situ* sample preparation may be limited by lack of knowledge of the biochemistry of any putative biomolecules, so it is possible that chemical incompatibilities could disrupt the structure of any molecules of interest. Additionally, “life as we don’t know it” may sustain cellular processes through another mechanism or structure. However, detection of ribosomes or ribosome-like structures with a solid-state nanopore instrument could support a hypothesis that samples are from a biologically active ecosystem, especially if both ribosomes and abundant molecules with RNA-like properties are identified.

Perhaps the most compelling astrobiological application of solid-state nanopore instrumentation is the search for signs of life in potentially habitable saline environments on Mars, Europa, or Enceladus. Salinity exerts a major effect on the ability of an organism or a biomolecule to persist in a system: microbial activity can be detected throughout wide ranges of salinity levels and salt compositions, and biological structures such as intact cells and biomolecules can be preserved even beyond the saline limits for active life (Klempay et al., 2021; Oren, 2008). Thus, many have postulated that saline environments should be a major target for astrobiological exploration, given their biological relevance and presence throughout the solar system (Davila and Schulze-Makuch, 2016; Neveu et al., 2020; Pappalardo et al., 2013; Phillips et al., 2023). Although saline environments are biologically relevant, they can also pose challenges for conducting science in such environments. Indeed, many analytical techniques are impeded by salt and require desalination treatments (Lawrence et al., 2023). Given SSN’s intrinsic requirement that samples be suspended in a conductive solution, it may be possible to use this technique to analyze environmental biomolecule samples within their original saline solutions or diluted with a high salt buffer of choice, without complex processing such as salt removal. Furthermore, Ocean Worlds are expected to be low in available free energy, and thus any existing biosphere would likely be low in biomass (Jones et al., 2018). SSN’s low limit of detection and high sensitivity makes it a promising technology for achieving the single-molecule resolution that would be necessary to effectively search for intact biomolecules in such environments.

Solid-state nanopores are of interest for future astrobiological space exploration due to their ability to detect biomolecules with high sensitivity and low false positive rates, as well as their potential utility in searching for agnostic biosignatures. Beyond its applications for astrobiological life detection, SSN has other qualities that overall make it well-suited for space payloads in general, including high portability, minimal resource requirements, low mass and power requirements, potential for analysis on minimally-prepared samples, and capability for rapid analysis. Interplanetary space missions with the explicit goal of searching for extraterrestrial life are expected in the coming decades (National Academies of Sciences, Engineering, and Medicine, 2022). With further research and development, a solid-state nanopore instrument may be a compelling addition to a future mission seeking to detect signs of extant life in the solar system.

## Acknowledgments

The authors thank Chen Chen for valuable contributions and comments on the machine learning components of this paper.

## Author contributions

J.M.M. and C.E.C. developed the concept; J.M.M. and C.E.C. performed the experiments; J.M.M. and C.E.C. analyzed the NanoCounter data; J.M.M., M.K., and C.E.C. performed the machine learning analysis. J.M.M. and C.E.C. wrote the paper. All authors edited and approved the paper. Biomolecule graphics (appearing in Figures 1, 3, 4) were created with BioRender.com.

## Data availability

Data can be downloaded at https://osf.io/56qkv/ and code for analysis can be downloaded at https://github.com/jmckaig/TAB.

