## Supplementary Data for "Translation as a Biosignature"

### Table of Contents

**Table S1.** Event and runtime data for various biomolecule samples in 2M LiCl…..……………S3

**Table S2.** Summary of various machine learning algorithms on SSN data………………………S4

**Figure S1.** Decision tree for SSN data.… ……………………………………………………..S5

**Table S2.** Summary of key features for Medium Tree classification……………………………..S6

**Table S1. Event and runtime data for various biomolecule samples in 2M LiCl.** Biomolecule runs are in green, while buffer runs are in white.

| Session ID | Sample type | Loaded biomolecule mass (ng) | Number of events | Run time (minutes) | Event rate (events/min) | Initial diam (nm) | Final diam (nm) |
|---|---|---|---|---|---|---|---|
| 1 | Startup buffer (buf1) | 0 | 3 | 5.38 | 0.56 | 41.91 | 41.94 |
| 1 | DNA plasmid pUC19 | 50 | 2284 | 6.62 | 344.86 | 41.97 | 41.48 |
| 1 | Startup buffer (buf2) | 0 | 6 | 6.35 | 0.94 | 40.89 | 40.69 |
| 1 | dsDNA 1kb Ladder | 2 | 919 | 5.93 | 155.04 | 40.84 | 40.18 |
| 1 | Startup buffer (buf3) | 0 | 1 | 5.73 | 0.17 | 40.22 | 40.22 |
| 1 | DNA Ladder High Range | 50 | 3115 | 5.99 | 520.00 | 41.30 | 40.33 |
| 1 | Startup buffer (buf4) | 0 | 1 | 5.60 | 0.18 | 40.60 | 40.60 |
| 2 | Startup buffer (buf5) | 0 | 7 | 3.18 | 2.20 | 32.24 | 31.78 |
| 2 | ssRNA ladder | 100 | 1874 | 10.03 | 186.85 | 23.57 | 23.02 |
| 2 | Startup buffer (buf6) | 0 | 110 | 9.15 | 12.03 | 30.28 | 30.10 |
| 2 | Startup buffer (buf7) | 0 | 74 | 2.55 | 29.02 | 30.63 | 29.65 |
| 2 | Startup buffer (buf8) | 0 | 123 | 4.08 | 30.16 | 30.35 | 29.41 |
| 2 | dsRNA ladder | 33.3 | 1398 | 11.01 | 127.02 | 28.49 | 27.35 |
| 2 | Startup buffer (buf9) | 0 | 641 | 3.23 | 198.76 | 25.30 | 24.66 |
| 2 | Startup buffer (buf10) | 0 | 191 | 3.19 | 59.94 | 28.87 | 28.84 |

| | | | | | | | |
|---|---|---|---|---|---|---|---|
| 2 | Startup buffer (buf11) | 0 | 186 | 3.06 | 60.73 | 23.85 | 23.58 |
| 2 | *E. coli ribosome* | 113.9 | 17077 | 10.03 | 1702.35 | 24.39 | 24.32 |
| 2 | Startup buffer (buf12) | 0 | 267 | 4.09 | 65.23 | 25.20 | 24.59 |
| 2 | Startup buffer (buf13) | 0 | 142 | 3.25 | 43.76 | 24.48 | 24.05 |

**Table S2. Summary of various machine learning algorithms on SSN data.** Table is sorted by testing accuracy. Model numbers are as indicated by the MATLAB Classification Learner app (MATLAB_R2023a).

| Model Number | Model Type | Model | Accuracy % (Validation) | Accuracy % (Test) |
|---|---|---|---|---|
| 3.29 | Neural Network | Wide Neural Network | 95.97 | 96.09 |
| 3.28 | Neural Network | Medium Neural Network | 96.14 | 95.85 |
| 3.30 | Neural Network | Bilayered Neural Network | 95.00 | 95.70 |
| 3.31 | Neural Network | Trilayered Neural Network | 95.42 | 95.66 |
| 3.11 | Support Vector Machine (SVM) | Quadratic SVM | 95.75 | 95.58 |
| 3.19 | K-Nearest Neighbors (KNN) | Cosine KNN | 94.80 | 95.54 |
| 3.21 | KNN | Weighted KNN | 95.06 | 95.42 |
| 3.27 | Neural Network | Narrow Neural Network | 95.53 | 95.30 |
| 3.23 | Ensemble | Bagged Trees | 95.60 | 95.26 |
| 3.20 | KNN | Cubic KNN | 94.91 | 95.18 |
| 3.13 | SVM | Fine Gaussian SVM | 95.03 | 95.10 |
| 3.14 | SVM | Medium Gaussian SVM | 94.88 | 95.10 |
| 3.17 | KNN | Medium KNN | 94.94 | 95.06 |
| 3.16 | KNN | Fine KNN | 94.41 | 95.02 |
| 3.1 | Tree | Fine Tree | 94.94 | 94.79 |
| 3.10 | SVM | Linear SVM | 94.78 | 94.55 |
| 3.18 | KNN | Coarse KNN | 93.98 | 94.23 |
| 3.22 | Ensemble | Boosted Trees | 94.56 | 93.72 |
| 3.26 | Ensemble | RUSBoosted Trees | 94.14 | 93.60 |
| 3.15 | SVM | Coarse Gaussian SVM | 92.02 | 93.60 |
| 3.25 | Ensemble | Subspace KNN | 92.78 | 93.13 |
| 3.7 | Efficient Linear SVM | Efficient Linear SVM | 92.23 | 92.50 |
| 3.32 | Kernel | SVM Kernel | 92.40 | 92.42 |
| 3.2 | Tree | Medium Tree | 92.89 | 92.34 |
| 3.33 | Kernel | Logistic Regression Kernel | 90.94 | 92.22 |
| 3.9 | Naive Bayes | Kernel Naïve Bayes | 91.38 | 90.36 |
| 3.5 | Discriminant | Quadratic Discriminant | 90.46 | 90.24 |
| 3.3 | Tree | Coarse Tree | 79.11 | 81.24 |
| 3.4 | Discriminant | Linear Discriminant | 78.67 | 78.95 |
| 3.8 | Naive Bayes | Gaussian Naïve Bayes | 70.69 | 70.02 |

| | | | | |
|---|---|---|---|---|
| 3.24 | Ensemble | Subspace Discriminant | 68.92 | 68.68 |
| 3.6 | Efficient Logistic Regression | Efficient Logistic Regression | 66.78 | 66.94 |
| 3.12 | SVM | Cubic SVM | 77.24 | 61.37 |

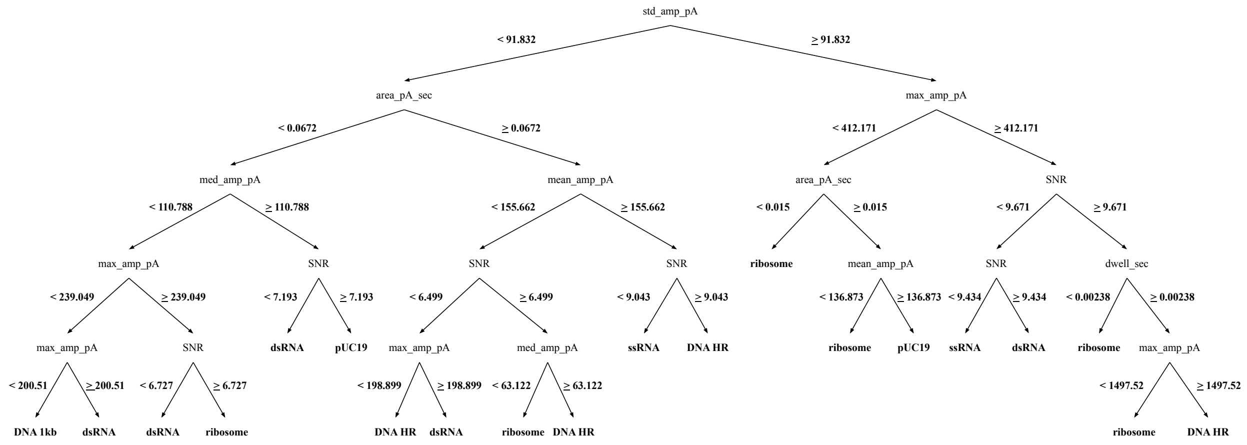


**Figure S1. Decision tree for SSN data.** This tree was generated by the Medium Tree model (3.2) by the MATLAB Classification Learner app (MATLAB_R2023a).

**Table S3. Summary of key features for Medium Tree classification.** Medium Tree is shown in Figure S1.

| **Feature** | **Description** |
|---|---|
| *dwell_sec* | Dwell time |
| *SNR* | Signal to noise ratio |
| *mean_amp_pA* | Mean current (in picoamps) |
| *max_amp_pA* | Maximum current (in picoamps) |
| *med_amp_pA* | Median current (in picoamps) |
| *std_amp_pA* | Standard deviation of current (in picoamps) |
| *area_pA_sec* | Area of current disruption (in picoamps) |
